# Infrared spectral markers for the nephroprotective effects of *Ficus deltoidea* in streptozotocin-induced diabetic rats

**DOI:** 10.1101/2020.12.23.424120

**Authors:** Nurdiana Samsulrizal, Goh Yong-Meng, Hafandi Ahmad, Nur Syimal’ain Azmi, Noor Syaffinaz Noor Mohamad Zin, Ebrahimi Mahdi

## Abstract

Fourier Transform Infrared (FTIR) is an established analytical technique to elucidate new discriminatory biomarkers. Our previous study showed that *Ficus deltoidea* (Ficus: Moraceae) is capable of increasing insulin secretion and improving tissue regeneration by reducing oxidative stress in diabetic rats. However, the assessment of treatment response is limited by the paucity of biomarkers. We aimed to evaluate the potential use of FTIR for assessing the nephroprotective effects of *Ficus deltoidea* (Ficus: Moraceae) in diabetic rats. A rat model of diabetes was induced using a single intraperitoneal injection of streptozotocin (STZ) (60 mg/kg body weight). Methanolic extract of *F. deltoidea* was administered orally at a dose of 1000 mg/kg body weight for eight weeks. Fasting blood glucose, serum insulin and kidney function parameters were examined. The kidneys were subsequently subjected to FTIR and histological analyses. Enzyme-linked immunosorbent assays (ELISA) assessed the levels of oxidative stress, antioxidant, and apoptosis-related proteins in the kidney tissue. The results show, for the first time, that there is a good agreement between changes in kidney and FTIR peaks. The IR peaks (1545 cm^−1^ and 1511 cm^-^1) corresponding to amide II were restored by treatment with *F. deltoidea*. Multivariate analysis demonstrated that the diabetic rats treated with *F. deltoidea* had similar clustering pattern that of the normal animals. Biochemical and histological examination further confirmed the nephroprotective effect of *F. deltoidea*. Thus, demonstrating how FTIR spectroscopy could be used for the diagnosis of diabetes kidney disease.

## Introduction

Diabetic nephropathy (DN) is a significant complication of long-standing hyperglycemia. It is characterized by proteinuria, glomerulosclerosis, thickening of the glomerular basement membrane (GBM), and tubulo-interstitial fibrosis [1]. DN has become an important health and economic encumbrance over the last five decades [2]. Animal and human studies consistently support hyperglycemia is the leading cause of end-stage kidney failure and accounts for 30-40 % of patients entering kidney transplant programs [3]. It has been reported that dialysis patients are associated with worrying life expectancy, comparable to or worse than that seen in many cancers [4]. Patients diagnosed with kidney complications had an average survival of only 5-7 years [5]. These data indicate that there are still aspects of diagnosis and pathogenesis that are needed for further investigations.

Poor glycemic control and accumulation of reactive oxygen species (ROS) play a significant role in the development of DN [6]. Glomerulus is particularly more vulnerable to ROS attack due to the presence of heparan sulfate proteoglycans, the anionic polysaccharides of the GBM [7,8]. However, recent studies have shown that all kidney cells, such as mesangial cells, podocytes and tubulointerstitial cells, are also liable to be affected by hyperglycemia [9].

Sustained kidney biochemical alterations have been observed in these kidney cells even after tight glycemic control [10]. According to previous reports [11, 12], there was a positive correlation between the structural and biochemical changes of the kidney tissue and the infrared (IR) absorption spectra. Therefore, the application of IR spectroscopic techniques in the study of diabetes kidney disease offers an excellent opportunity to improve diagnosis.

FTIR spectroscopy is a simple, label free, non-invasive, and highly reproducible analytical technique. Application of FTIR spectroscopy combined with chemometrics identifies the specific molecular fingerprint of biological samples, which in turn provides clues to the biochemical and pathological changes. FTIR spectroscopy supported with quantitative and qualitative spectra analysis have been shown possible to discriminate between benign and cancer tissue [13,14]. Several earlier studies have also demonstrated the advantages of FTIR spectroscopy as a non-invasive technique for elucidating the chemical features of cells and tissues [15-18], in part attributed to chronic complications of diabetes. The success in application of FTIR spectroscopy for the detection of glucose in serum [19,20], whole blood [21] and urine samples [22] have previously been reported. FTIR spectroscopy has been successfully used for monitoring the apoptotic cell death [23] and antioxidant activity [24]. Our previous studies demonstrated that FTIR spectroscopy could be used to provide a systemic snapshot and relevant to monitor the pancreas and brain pathological changes of diabetic rats [25,26]. However, analysis of the structure and biochemical changes of the diabetic kidney disease by FTIR remains elusive, and this is where the novelty of the current research lies.

*F. deltoidea* (Moraceae) is an evergreen shrub, or small tree that is easily found in Malaysia and widely distributed in Southeast Asian countries such as Thailand, Sumatra, Java, Kalimantan, Sulawesi, and Moluccas. The decoction of *F. deltoidea* has traditionally been used in postpartum care specifically to improve uterine strength, regain energy, and prevent postpartum bleeding [27,28]. It is also served as a health tonic or taken as herbal tea to relieve headache, fever, and toothache [29]. Acute toxicity studies showed that the median lethal dose (LD_50_) of aqueous and ethanolic leaf extracts of *F. deltoidea* was greater than 5000 mg/kg bwt [30] and 2000 mg/kg bwt [31], respectively. Meanwhile, Abrahim et al. [32] showed that *F. deltoidea* leaves did not exhibit any cytotoxic effects on the normal liver cells. *F. deltoidea* is known to have a glucose-lowering effect and exhibit antioxidant activity [33, 34]. Our previous work showed that treatment with *F. deltoidea* decreased blood glucose, increased insulin secretion, and enhanced pancreatic islet regeneration in STZ-induced diabetic rats [25]. We also found *F. deltoidea* possesses neuroprotective properties, leading to the improvement of spatial learning and memory and cortical gyrification patterns [26]. However, the potential of using FTIR for monitoring the structural and biochemical changes in the kidney of diabetic rats following *F. deltoidea* treatment remains elusive. We hypothesize that the non-destructive technique FTIR spectroscopy in combination with multivariate analyses would be able to identify the spectral markers corresponding to the biochemical and pathological changes in the kidney of diabetic rats.

## Materials and Methods

### Plant material and extract preparation

The leaves of *F. deltoidea* var. *kunstleri* were purchased from Moro Seri Utama Enterprise, Batu Pahat, Johor, Malaysia. The plant material was identified and authenticated by a specialized taxonomist. A voucher specimen (UKMB-40315) was deposited in the Herbarium Unit, Universiti Kebangsaan Malaysia for further reference. The leaves were washed thoroughly, oven-dried at 37 ± 5°C, ground to a fine powder in an electric grinder, and weighed. The powdered leaves (100 g) were soaked in 1 L absolute methanol for three days at room temperature. Liquid extracts were concentrated using a rotary vacuum evaporator (R-215, Buchi, Switzerland) under reduced pressure. The extracts were kept in tightly closed glass containers and stored at −20°C until further use.

### Animals

Male Sprague-Dawley rats weighing around 100 – 200 g were purchased from Chennur Suppliers, Malaysia. The animals were housed in polypropylene cages (47 × 34 × 20 cm) and maintained under standard laboratory conditions with a 12 h light/dark cycle, at a temperature of 25 ± 1 °C, throughout the study. Animals were fed with standard rat chow (Gold Coin Holdings, Kuala Lumpur, Malaysia) and tap water *ad libitum* before and during the experiments. The animals were anaesthetized with ketamine (80 mg/kg) and xylazine (8 mg/kg), followed by terminal exsanguinations. All study protocols, including DN induction and sacrifice operation were approved by the ethics review board of Universiti Putra Malaysia, Animal Care and Use Committee (UPM/IACUC/AUP-R090/2014).

### Animal models of DN

DN was induced experimentally in 12-h fasting rats except for the normal group through a single intraperitoneal injection of 0.5 mL STZ (Sigma-Aldrich, Deisenhofen, Germany) at a dosage of 60 mg/kg of body weight. STZ in rats is a reproducible animal model of progressive tubular damage and glomerular injury, presenting histological features that resemble human DN [35]. Seven days after STZ treatment, the fasting blood glucose levels were determined using a blood glucose meter (Accu-Chek Performa, Roche, Germany).

Animals with fasting blood glucose levels above 11 mmol/L for three consecutive days were considered diabetic, while those outside this range were excluded [36].

### Experimental design

Twenty-two rats were randomly divided into four groups (n=6 per group) as follow: Group normal control (NC): Rats were administered with 0.5 mL saline.

Group diabetic control (DC): The diabetic rats were treated with 0.5 mL saline.

Group metformin-treated diabetic rats (DMET): The diabetic rats were treated with 1000 mg/kg of metformin.

Group *F. deltoidea*-treated diabetic rats (DFD): The diabetic rats were treated with 1000 mg/kg of *F*.*deltoidea*.

The LD_50_, of metformin and *F*.*deltoidea* in rodents had been reported to be greater than 1500 mg/kg/day [37] and 5000 mg/kg/day [30], respectively. All treatments were given once daily via oral gavage for eight weeks. The fasting blood glucose was measured at one-week intervals by pricking the sharp needle from the tail. Animals were restrained using a plastic container while pricking the tail. The restrain is used to keep the animals comfortable and safe as well as prevention of unexpected movements during manipulation under conscious condition.

### Experimental procedure

At the end of the experiment, all animals were fasted overnight to reduce variability in basal blood glucose [38]. The terminal urine was collected for 4 hours immediately preceding sacrifice in individual metabolic cages. Blood samples (10-15 mL) were collected via cardiac puncture from the rats, centrifuged at 4000 *g* for 15 min, and serum was stored in aliquots at − 80°C. Half of each sample was kept in 10% formalin for tissue characterization, and the other half was stored at −80°C prior for biochemical analysis. Serum creatinine, urea, uric acid and total bilirubin were determined by the auto analyser (Hitachi 911, Boehringer-Mannheim, Germany).

### Glomerular filtration rate

Glomerular filtration rate (GFR) based on creatinine clearance was determined using urine and serum creatinine assay kits (Cayman, MI, USA) as well as urine output levels. GFR was calculated using the following formula:

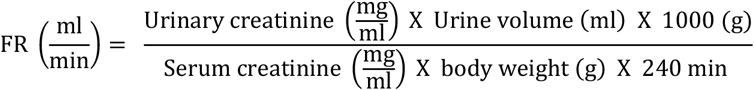

### Determination of spot urine creatinine

The spot urine creatinine (UCRE) was determined quantitatively using enzyme-linked immunosorbent assays (Cayman, MI, USA). ELISA tests were conducted in triplicate and performed according to the manufacturer protocol.

### Preparation of Tissue Homogenates

One hundred milligram of kidney tissue was homogenized in a homogenizing buffer (50 mM Tris-HCl, 1.15% KCl pH 7.4) using a Teflon pestle (Glass-Col, USA) at 900 rpm. The homogenates were centrifuged at 9,000 *g* in a refrigerated centrifuge (4°C) for 10 min to remove nuclei and debris. The supernatant obtained was used for FTIR analysis and biochemical assays. Protein concentration was estimated by the method of [39], using bovine serum albumin as the standard.

### FTIR analysis

FTIR measurements were performed to analyze the structural and biochemical properties of the kidney rats. The FTIR spectra were recorded using a Bruker 66V FTIR spectrometer (Bruker Corp., MA, USA). Twenty microliter of kidney homogenate was deposited on a liquid cell (demounted cell) using a pipette according to the method reported by Demir et al. [40]. All spectra were recorded over the range 4000-400 cm^−1^ frequency range versus a clean prism background at room temperature. Three spectra were recorded for each treatment group. Spectra analysis was performed using OPUS 7.0 software (Bruker Optics, GmbH).

All the spectra were pre-processed, and second derivative spectra were calculated using Savitzky-Golay algorithm. Second derivative spectra were taken using a 13-point smoothing factor to resolve differences in peak intensities among experimental animals. Second derivative spectra were further used to quantify the features of interest by integration and statistical assessment (Origin Pro 9.1). The fingerprint region from 1800-800 cm^−1^ was used for all subsequent data analysis. The prediction of IR spectral was performed based on the peak intensities obtained from the integrated peak area of the selected frequency limits.

### Chemometric analyses

All the registered spectra were subjected to multivariate analysis (MVA), including hierarchical cluster analysis (HCA) and principal component analysis (PCA). All analyses will be conducted using the OriginPro software (OriginLab, Northampton, MA, USA). The unsupervised analysis of PCA and HCA will be performed for general overview of clustering patterns.

### Estimations of apoptosis-related proteins levels

The levels of Casp 8, JNK, Casp 9, p53 and Bcl-2 in the kidney supernatant were measured by a double-antibody sandwich ELISA, using a commercially available kit from Qayee-Bio (Shanghai, China) according to the manufacturer’s instructions. The absorbance was determined spectrophotometrically at a wavelength of 450 nm using a spectrophotometer (Epoch 2 microplate spectrophotometer, BioTek Instruments, Inc, Vermont, USA).

### Biochemical estimations in kidney homogenates

Kidney oxidative stress biomarkers such as GPX, SOD and MDA levels were determined using commercially available assay kits (Cayman, MI, USA). The absorbance change vs time for each biomarker was measured spectrophotometrically at a specific wavelength using a spectrophotometer (Epoch 2 microplate spectrophotometer, BioTek Instruments, Inc, Vermont, USA).

### Histological assessment

The kidney was fixed in 4% formaldehyde in phosphate buffered saline (Invitrogen, Carlsbad, CA), embedded in paraffin blocks, sectioned at 4 µm, and stained with H&E. All slides were examined using light microscopy (Motic BA410, Wetzlar, Germany) equipped with a digital camera (Moticam Pro 285A, Wetzlar, Germany) under a magnification of 400.

### Statistical analysis

Results were presented as the mean ± 1 SD and dissected using the Statistical Package for the Social Sciences (SPSS) 26 software (IBM Corporation, USA). The differences between the groups were tested using one-way ANOVA followed by Duncan’s multiple range test. All analysis was performed at 95% confidence level.

## Results

### Spectral markers corresponding to structural and biochemical changes in the kidneys of DN rats

Fig 1(a) shows the kidney FTIR absorption of the experimental groups in the regions of 4000-400 cm^−1^. The most obvious differences between the absorption spectra were found in the fingerprint region of 1800-800 cm^−1^. Fig 1(b) represents the second derivative of the kidney FTIR spectra for the selected spectral regions presented in Fig 1(a). Six potential peaks of interest (1545 cm^−1^, 1511 cm^−1^, 1400 cm^−1^, 1246 cm^−1^, 1181 cm^−1^ and 1143 cm^−1^) were identified by comparing the average second derivative spectra of each group. The most important peaks for describing the variations between the experimental animal groups are marked in Fig 1(c). An absorption band at 1545 cm^−1^ and 1511 cm^−1^ are related to amide II functional groups of proteins [41]. The intense band at 1400 cm^−1^ arises from the methyl and methylene groups from lipids and proteins [41] whereas the band at 1246 cm^−1^ corresponds to the absorption of DNA/RNA phosphate backbones [43]. Other bands at 1181 cm^−1^ and 1143 cm^−1^, respectively, are related to C–O vibrations from glycogen and other carbohydrates [44].

**Fig 1.**
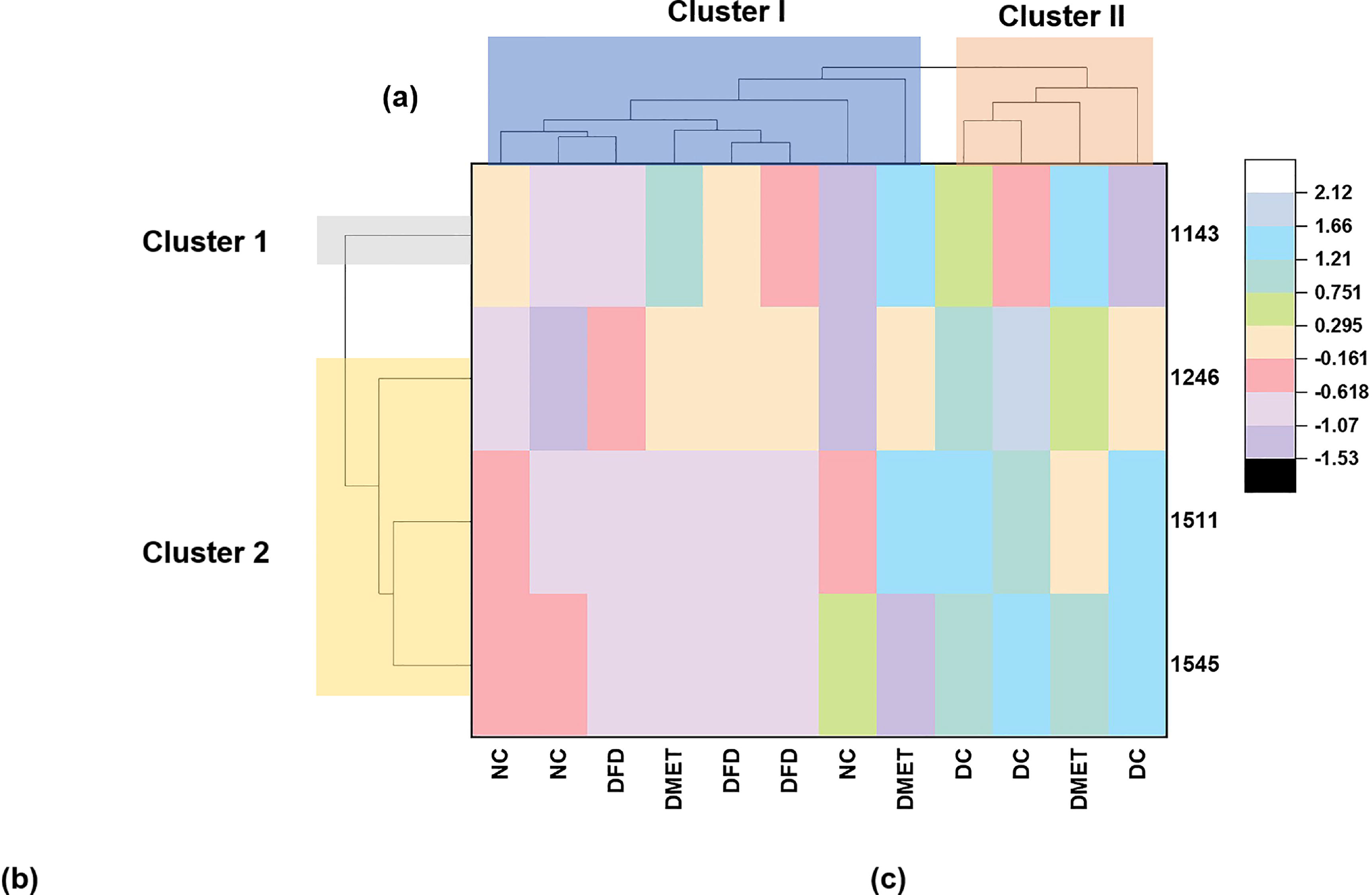
FTIR analysis of the rat kidney. (a) The average FTIR absorption spectra of kidney tissue in the regions of 4000-400 cm-1 (upper graph) and 1800-800 cm^−1^ (lower graph). (b) The average second derivative spectra in the range of 1800-800 cm^−1^ acquired from the kidney of the treatment groups. (c) Peaks of interest potentially relevant to structural and biochemical changes in the rat kidney. The analysis of these spectra was performed on n = 3 animals for each group.

We proceeded to compare the peak intensities obtained from the integrated peak area of the selected frequency limits. Table 1 showed the peak intensity of the six IR features. The intensity of the IR spectra was only significantly different at 1545 cm^−1^, 1511 cm^−1^, 1246 cm^- 1^, and 1143 cm^−1^. The intensity of amide II peaks (1545 cm^−1^, 1511 cm^−1^) was increased in the DC animals, indicating an increase of β-peptides [46]. However, the intensity of peaks decreased significantly following treatment with metformin and *F. deltoidea*. The intensity of peak at 1246 cm^−1^, which corresponds to the PO_2-_ and C-O absorption of DNA/RNA polysaccharide backbones, remained higher in the kidney of DC, DMET and DFD rats as compared to the NC group. The DMET group had a significantly higher peak intensity at 1143 cm^−1^ than other groups. These results suggest that the peak intensity of FTIR spectra could be relevant to estimating treatment effects from observational data.

**Table 1.** The intensity of potential IR marker bands in the kidney of experimental groups.

| Peak parameters (cm<sup>-1</sup>) | Frequency limits for integration (cm<sup>-1</sup>) | Groups |  |  |  |
| --- | --- | --- | --- | --- | --- |
|  |  | NC | DC | DMET | DFD |
| 1545 | <sup>1</sup>1548 - 1539 (IPA x 10<sup>-3</sup>) | 1.29 ± 0.15<sup>a</sup> | 1.88 ± 0.06<sup>b</sup> | 1.09 ± 0.31<sup>a</sup> | 0.90 ± 0.05<sup>a</sup> |
| 1511 | <sup>2</sup>1514 - 1504 (log<sub>10</sub> IPA) | -2.80 ± 0.04<sup>a</sup> | -2.41 ± 0.03<sup>b</sup> | -2.64 ± 0.13<sup>a</sup> | -2.87 ± 0.03<sup>a</sup> |
| 1400 | 1412 - 1392 (IPA x 10<sup>-3</sup>) | 2.99 ± 0.91 | 3.65 ± 1.21 | 5.85 ± 1.97 | 3.57 ± 1.06 |
| 1246 | 1262 - 1239 (IPA x 10<sup>-3</sup>) | 0.69 ± 0.08<sup>a</sup> | 1.72 ± 0.27<sup>b</sup> | 1.35 ± 0.10<sup>b</sup> | 1.20 ± 0.07<sup>b</sup> |
| 1181 | 1186 - 1175 (IPA x 10<sup>-3</sup>) | 0.59 ± 0.10 | 0.77 ± 0.10 | 0.70 ± 0.06 | 0.72 ± 0.11 |
| 1143 | 1148 – 1138 (IPA x 10<sup>-3</sup>) | 0.35 ± 0.07<sup>a</sup> | 0.39 ± 0.12<sup>a</sup> | 0.79 ± 0.03<sup>b</sup> | 0.42 ± 0.07<sup>a</sup> |

All data were subjected to normality Ryan-Joiner (similar to Shapiro-Wilk) and Kolmogorov-Smirnov testing. The integrated peak area (IPA) from frequencies <sup>1,2</sup> were log<sub>10</sub> transformed to achieve normal data distribution (*p*-value of normality testing > 0.05). The data were expressed as mean ± SEM and analyzed by one-way ANOVA followed by Duncan post hoc test. Superscripts <sup>a,b</sup> represents significant differences of IPA among the experimental groups in each of the respective frequencies.

Two-way hierarchical clustering heat-map analysis (HCA) was subsequently used to discriminate the extracted dataset filtered by ANOVA (P < 0.05) into subgroups according to their similarities. As illustrated in Fig 2, the row tree represents the IR features, the column tree corresponds to the treatment groups, and the colors in the heat table represent the intensities of the IR features dataset. HCA stratified the experimental animals into two major clusters that differed by the relative degree of IR intensities. Cluster I influenced by the NC and DFD groups, while Cluster II seemed to be more related to the DC group. Two animals from the DMET groups were put into Cluster I whereas another one was clumped in Cluster II. The IR features were further discriminated into two clusters (Cluster 1 and 2), each of which revealed the intensity distributions in different experimental animals. The peak 1545 cm^−1^, 1511 cm^−1^ were least abundant in the NC and DFD groups. However, increased variation across the experimental animals was observed in the spectra region at 1143 cm^−1^, leading to a separate cluster in the dendrogram.

**Fig 2.**
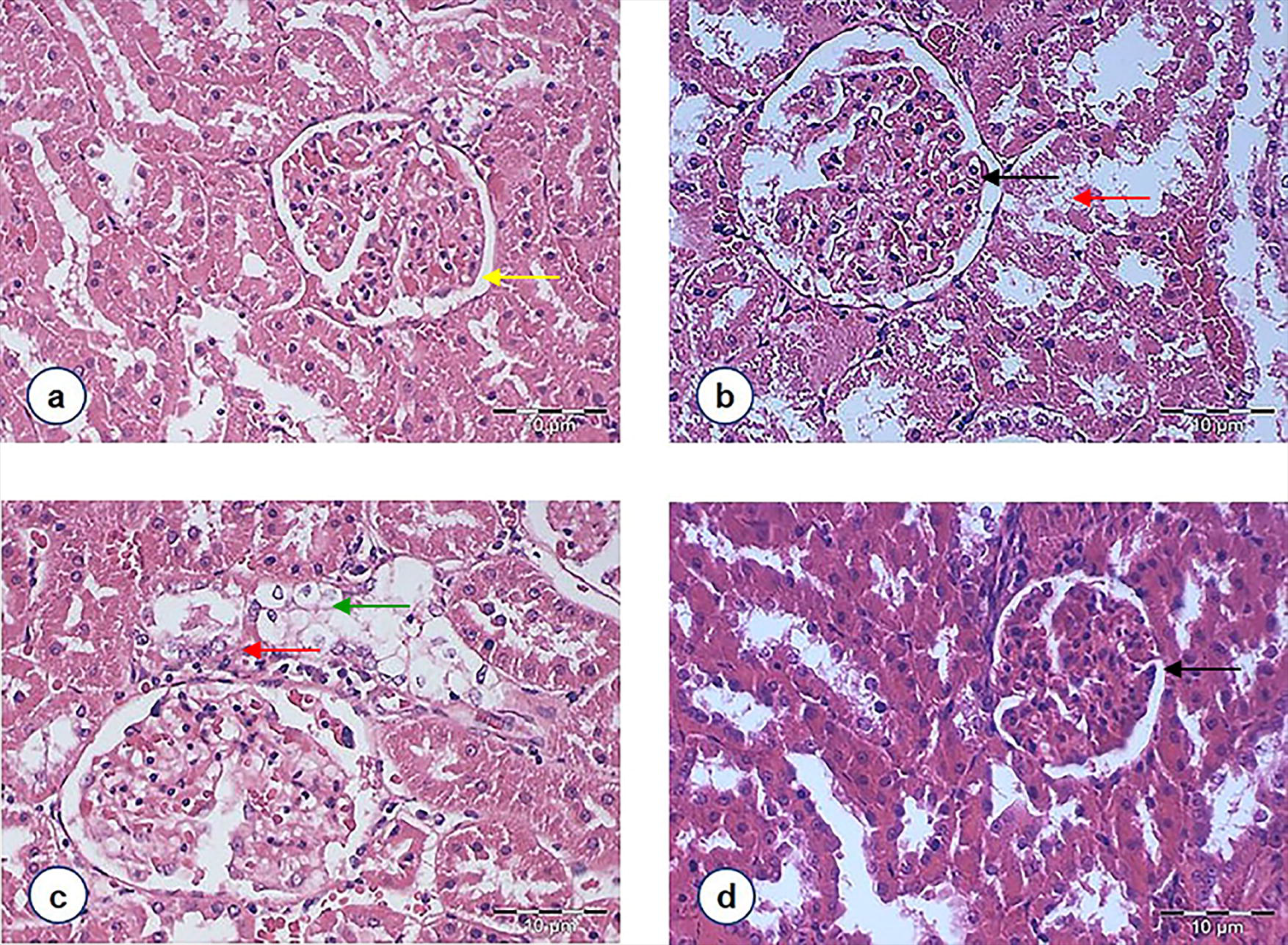
Two-way hierarchical clustering heat-map of kidney IR intensities. Each column shows the peak intensities of individual animals from the NC, DC, DMET and DFD groups (n = 3). The amount of each peak intensity is expressed as the total integrated area and is represented by the color scheme. A white and black indicate high and low concentrations of peak intensity, respectively.

Principal component analysis (PCA) was performed to verify the empirical clusters produced by HCA Fig 3(a) shows the PCA biplot obtained for the first two principal components after PCA with the second derivative spectra of IR features. The first principal component (PC1) explains 52.52% of the variation between the four spectroscopic changes, and the second principal component (PC2) accounts a further 32.57% of the variation. Together, those two axes explain 85.09% of total variance. This analysis also explained that the DC group was separated from the NC and DFD groups by PC1, indicating their different metabolomic profiles. The NC and DFD groups could be further separated by PC2. The loading plot showed that PC1 correlates with the 1545 cm^−1^, 1511 cm^−1^ and 1246 cm^−1^ whereas the spectral features at 1143 cm^−1^ were positively contributed to the PC2. The difference between the experimental groups was further visualized by a three-dimensional (3D) PCA analysis on PC1 [Fig 3(b)]. The 3D plot of PC1 allows comprehensive visualization and separation of the experimental groups as the variables are highly correlated with PC1 than with PC2. The kidneys of DC were separated from other groups along the PC1. The similarity of HCA and PCA results emphasize the underlying distributions within this dataset. Our data thus suggest that the wavenumbers identified in the fingerprint region can discriminate the nephroprotective effect of treatments with high accuracy.

**Fig 3.(a) Principle component analysis (PCA) analysis represents the scores of samples (dots) and loading of variables (vectors) of second derivative spectra of kidney tissue on the first two principal components (85.09% of total variance). (b) 3D biplot of PC1 displaying the separation of NC, DC, DMET and DFD groups.**

### *F. deltoidea* reduced fasting blood glucose, increased serum insulin and improved biochemical parameters of DN rats

After 56 days of the experiment, animals in the DC group exhibited significant increases in fasting blood glucose and serum creatinine levels as compared to the NC group. The diabetic rats also showed a significant decrease in serum insulin, urinary creatinine, total serum bilirubin and glomerular filtration rate (Table 2). However, the level of fasting blood glucose decreased, and serum insulin levels and glomerular filtration rate increased in the DFD group. The serum creatinine and total bilirubin also returned to near normal levels in response to *F. deltoidea* treatment.

**Table 2.** Effect of *F. deltoidea* on fasting blood glucose, insulin and kidney function markers and glomerular filtration rate of the experimental groups.

| Parameter | NC | DC | DM | DFD |
| --- | --- | --- | --- | --- |
| Fasting blood glucose (mmol/L) | 4.93 ± 0.21<sup>a</sup> | 30.13 ± 2.63<sup>b</sup> | 19.83 ± 3.75<sup>c</sup> | 17.27 ± 4.97<sup>c</sup> |
| Serum insulin | 4.16 ± 3.03<sup>c</sup> | 1.58 ± 0.16<sup>a</sup> | 1.78 ± 0.34<sup>a</sup> | 2.41 ± 0.08<sup>b</sup> |
| Serum creatinine (μmol/L) | 64.50 ± 2.12<sup>a</sup> | 92.50 ± 3.54<sup>b</sup> | 66.00 ± 1.41<sup>a</sup> | 69.50 ± 2.12<sup>a</sup> |
| Urinary creatinine (mg/dL) | 94.09 ± 3.27<sup>c</sup> | 6.81 ± 0.18<sup>a</sup> | 9.77 ± 0.36<sup>a</sup> | 39.59 ± 9.82<sup>b</sup> |
| Urea (mmol/L) | 9.37 ± 0.97 | 12.83 ± 1.00 | 15.80 ± 4.31 | 10.37 ± 1.70 |
| Uric acid (μmol/L) | 213.65 ± 7.43<sup>a</sup> | 241.90 ± 49.22<sup>a</sup> | 266.30 ± 5.94<sup>a</sup> | 183.80 ± 30.55<sup>a</sup> |
| Total bilirubin (μmol/L) | 2.00 ± 0.130<sup>b</sup> | 1.00 ± 0.28<sup>a</sup> | 1.15 ± 0.21<sup>a</sup> | 2.40 ± 0.14<sup>b</sup> |
| Glomerular filtration rate (mL/min) | 2.62 ± 0.19<sup>b</sup> | 1.34 ± 0.28<sup>a</sup> | 1.60 ± 0.36<sup>a</sup> | 4.32 ± 0.52<sup>c</sup> |

Values are mean ± 1SD for six rats in each group. Values with different superscripts down the column indicate significant difference at p<0.05.

### *F. deltoidea* decreased apoptosis-protein levels in the kidneys of DN rats

The levels of caspase 8 and caspase 9 were significantly increased in the kidney of DC rats compared to the NC group (Table 3). However, the DFD animals had a significant decrease in kidney caspase 8 levels.

**Table 3.** Apoptosis-related proteins in the kidney tissues of the experimental groups.

| Groups | Casp 8 (pg/ml) | JNK (pg/ml) | Bcl2 (ng/ml) | Casp 9 (pg/ml) | p53 (ng/ml) |
| --- | --- | --- | --- | --- | --- |
| NC | 376.30 ± 91.15<sup>a</sup> | 1300.40 ± 152.13 | 526.88 ± 11.25<sup>ab</sup> | 1007.97 ± 184.89<sup>a</sup> | 73.20 ± 20.98 |
| DC | 614.44 ± 19.28<sup>c</sup> | 1107.54 ± 231.45 | 541.88 ± 9.44<sup>b</sup> | 1368.98 ± 223.68<sup>b</sup> | 64.98 ± 11.07 |
| DMET | 515.19 ± 24.25<sup>b</sup> | 1190.87 ± 108.60 | 507.71 ± 54.78<sup>a</sup> | 1299.54 ± 44.13<sup>b</sup> | 57.85 ± 5.49 |
| DFD | 496.67 ± 13.65<sup>b</sup> | 1069.44 ± 56.66 | 561.04 ± 17.25<sup>ab</sup> | 1260.65 ± 51.62<sup>ab</sup> | 66.21 ± 7.68 |

Data are presented as mean ± 1 SD for six rats in each group. Values with different superscripts in a row differed significantly at p<0.05.

### *F. deltoidea* decreased malondialdehyde and improved antioxidants in the kidneys of DN rats

Table 4 shows that the levels of MDA and SOD were significantly increased, whereas GPx activity was significantly decreased in the kidney of the DC group. In contrast, treatment with *F. deltoidea* resulted in a significant increase in renal GPx and SOD activities, as well as a significant reduction in MDA values.

**Table 4.**
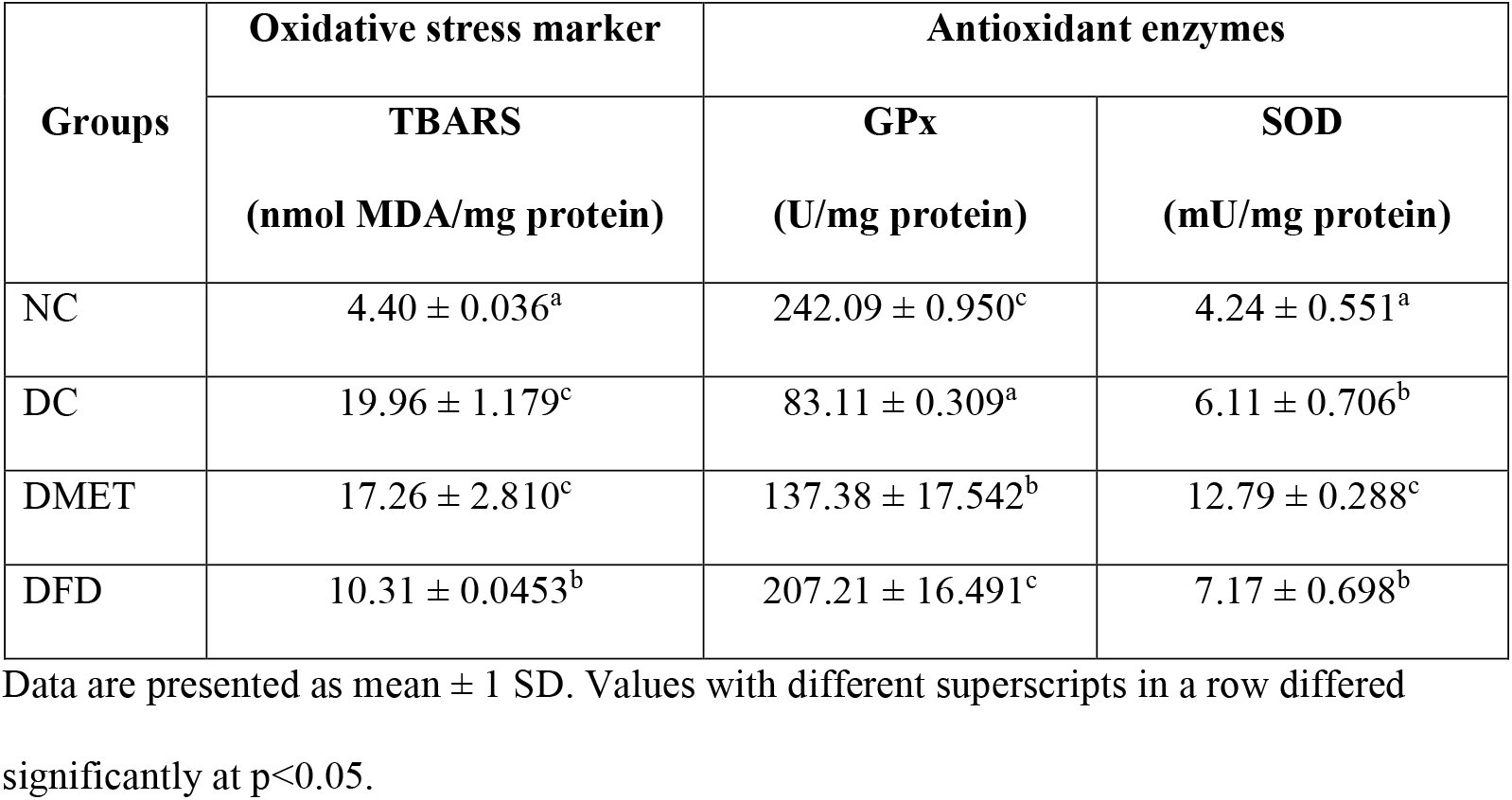
Oxidative stress marker and antioxidant enzymes of various experimental groups.

### *F. deltoidea* alleviated structural changes in the kidneys of DN rats

A histological examination of kidney sections was conducted to confirm the nephroprotective effects. Light microscopic examination revealed normal cyto-architecture of the kidney in the NC rats [Fig 4(a)]. However, the kidney structure of DC rats showed marked morphological irregularities. Capillary loops are poorly defined consistent with wider capsular spaces as well as shrinkage hypercellular congested glomeruli. Severe tubular degeneration and tubular necrosis were also noted [Fig 4(b)]. It was observed that the DMET animals had equally severe changes in the proximal convoluted tubules, which consisted of cytoplasmic vacuolation of tubular epithelial cells and cellular swelling [Fig 4(c)]. Although some irregular capillaries, in part attached to Bowman’s capsule, were noted among the DFD group [Fig 4(d)], the degree of necrosis of the tubular epithelium was nevertheless lessened by *F. deltoidea* extract.

**Figure 4. Light photomicrographs of glomeruli and renal tubules in the kidney of rats from different experimental groups (magnification 400X, HE staining):** (a) Normal control rats showing normal glomerulus with parietal cells line the external surface of Bowman space [indicated by the yellow arrow]. (b) STZ-induced diabetic rats were depicting a glomerulus with irregular capillaries [indicated by the black arrow] and tubular necrosis [indicated by red arrow]. (c) Diabetic rats treated with metformin showing that the lumina of the tubules are closed by swollen epithelial cells [indicated by the green arrow]. (d) Diabetic rats treated with *F. deltoidea* revealed that the degree of necrosis of the tubular epithelium was lessened by *F. deltoidea* extract, but capillary loops remain poorly defined [indicated by the black arrow].

## Discussion

FTIR spectroscopy was applied the first time to identify spectral markers associated with the nephroprotective effects of *F. deltoidea* in diabetic rats. We demonstrated that biochemical and structural changes in the kidney of rats would leave a signature in the FTIR spectra. Significant differences in the peak intensity at 1545 cm^−1^, 1511 cm^−1^ and 1246 cm^−1^ were consistent with observations of increased caspase 8 and MDA levels in the kidney of diabetic animals that had survived with compromised kidney function and morphological irregularity. Increased serum creatinine levels as well as decreased GFR, urinary creatinine total bilirubin, and GPx further strengthens the association between the IR peak intensity and the existence of kidney failure in STZ-induced diabetic rats.

Second derivative FTIR spectra of the DC animals displayed an intense sharp signal at 1545 cm^−1^ and 1511 cm^−1^ (Fig 1). A significant intensity difference between treatment groups was also observed at these peaks, which are corresponding to amide II functional groups (Table 5). These findings were consistent with a study showing that the β-sheet content increased in parallel with the glucose concentration levels [45]. An increase in the FTIR peak intensity usually indicates an increase in the amount of the functional group [46]. A study done by Ye et al. [47] demonstrated that FTIR measurement variations in the amide II bands are related to proteins aggregation. Higher protein aggregation has been described to positively correlate with intracellular β-sheet contents and apoptosis [48], implying that an increase in amide II peak intensity results from an increased sensitivity of kidney cells to apoptosis. However, some works have reported the nephroprotective role of amide in animal models [49]. It is important to note that apoptosis is not the only form of programmed cell death, but it is required for kidney cell proliferation and tissue regeneration after injury [50]. Previous findings have shown that the bands related to amide II indeed were more intense in the differentiated kidney cell [12]. Therefore, it is conceivable to suggest that the IR peak of amide II could be a potential spectral marker to indicate the kidney apoptosis index.

HCA and PCA analyses demonstrated that the treatment effects could also be distinguished at 1246 cm^−1^, which is dominated by the DNA phosphate backbones (Figs 2 and 3). As shown in Table 5, both untreated and treated diabetic rats have a significantly higher peak intensity at 1246 cm^−1^ as compared to the NC groups. A similar observation has been described for pulmonary hypertension rat model [51]. Sofińska et al. reported that increasing intensity of phosphate vibration indicates DNA fragmentation [52]. High kidney DNA fragmentation has indeed been demonstrated in STZ-induced diabetic rats [53]. Oxidative stress is considered causal of DNA fragmentation in chronic kidney disease [54]. An increase in DNA fragmentation usually correlates with the apoptotic progress [55]. This is consistent with our current observation that the 1545 cm^−1^ and 1511 cm^−1^ bands are directly proportional to band at 1246 cm^−1^. Another study has nevertheless shown that the apoptotic index is inversely proportional with the spectral area 900-1250 cm^−1^ [56]. However, the study is limited to human HL60 leukemic cells. Therefore, measuring the levels of apoptosis-related proteins and lipid peroxidation in the kidney tissues are essential to support the hypothesis that changes in FTIR spectra absorption bands are reliable to indicate biochemical changes in the kidney of rats.

GFR is a key indicator of kidney function [57]. We showed that the GFR decreased following STZ injection, which is inversely proportional to the peak intensity at 1545 cm^−1^, 1511 cm^−1^ and 1246 cm^−1^ (Table 2). This observation is consistent with previous reports that long-standing hyperglycemia causes GFR decline [58, 59], and an increase in the amide II and DNA related regions were associated with chronic kidney diseases [13, 60]. Interestingly, the DFD animals with lower peak intensity at 1545 cm^−1^, 1511 cm^−1^ and 1246 cm^−1^ had significant increases in the GFR over the treatment period. This result is of particular importance because an increase in the GFR implies delayed progression of chronic kidney disease [61, 62] and adds further evidence demonstrating the potential of FTIR spectroscopy to estimate kidney function.

The role of apoptosis in kidney cell injury has been reported in several studies [63,64]. To further investigate the association between the IR spectral changes and the degree of kidney injury, the levels of apoptosis-related proteins in kidney tissue were measured by ELISA. As shown in Table 3, higher values of caspase-8 and caspase-9 were observed in the kidney of DC rats. A similar finding was reported by Pal et al. [65]. The kidney caspase-8 and caspase-9 levels were nevertheless decreased in the DFD and DMET rats that had lower peak intensity at 1545 cm^−1^, 1511 cm^−1^ and 1246 cm^−1^ compared to the DC group. There is compelling evidence supporting the link between caspase-8 and β-sheet structure in the apoptotic cell death [66,67]. It has been suggested that the cleavage of protein may contribute to the β-sheet structure, which in turn induces apoptotic cell death by targeting mitochondria. In parallel with FTIR spectra and histological findings, we confirm the nephroprotective effects of *F. deltoidea*, thus supporting our hypothesis that the peak appeared at 1545 cm^−1^ and 1511 cm^−1^ could be a reliable spectral marker to indicate the degree of kidney injury.

We showed that MDA levels were increased and the activities of GPx decreased in the kidney of DC rats similar to that reported in previous studies [68]. However, the elevation of kidney SOD level contradicted previous studies. These findings suggest that STZ promotes kidney injury by increasing intracellular hydrogen peroxide. Hydrogen peroxide (H_2_O_2_) has been reported to potentiate evoked calcium influx in the podocytes, an important role in the development of kidney damage [69]. It has also been demonstrated that H_2_O_2_ contributes to kidney cellular injury and necrosis by modulating inflammatory mediators [70,71].

Therefore, the balance of SOD and GPx is more important to counteract the increase in oxidative stress induced by STZ than the level of SOD alone [72]. Intriguingly, *F. deltoidea* treatment resulted in the elevation of GPx to SOD ratio as well as a significant reduction in MDA levels. DNA fragmentation has been reported to positively correlate to MDA levels [73]. These results indeed demonstrated that animals with lower kidney MDA levels displayed a lower peak intensity at 1246 cm^−1^ than the DC group, evidencing the differences in absorbance intensities recorded by FTIR spectroscopy can be used to quantify oxidative DNA damage.

Histological analysis of the kidney demonstrates STZ caused early onset of glomerulosclerosis, severe tubular degeneration, and tubular necrosis at the cortico-medullary zone (Fig 3B). In agreement with these observations, reported the presence of glomerulosclerosis in STZ-induced rat model of diabetes [74]. Glomerulosclerosis is commonly characterized by distortion of capillary architecture, that is, the leading cause of kidney failure [75]. Although capillary loops remain poorly defined in all treated groups, the degree of necrosis of the tubular epithelium was reduced by *F. deltoidea* treatment (Fig 3D). These histological features were consistent with the findings of FTIR results shown in Fig. 1 and Table 1. Notably, the DFD group that had shown marked improvement in kidney morphology are associated with a significant increase in urinary creatinine, serum total bilirubin and peak intensity at 1545 cm^−1^, 1511 cm^−1^ and 1246 cm^−1^. The serum creatinine decreased to near average level. It is worth mentioning that an increase in serum bilirubin levels and decreased serum CREA usually indicates a nephroprotective effect [76-78]. Meanwhile, the elevation of bilirubin protects the kidney from systemic oxidative stress in the vascular compartment [79]. It is, therefore, reasonable to suggest that *F. deltoidea* possesses nephroprotective activity in STZ-induced diabetic rats and FTIR may allow for the detection of treatment effect on the kidney earlier than is morphologically evident.

## Conclusion

We suggest that the application of FTIR combined with chemometrics can be used to identify biochemical alterations that precede morphological and functional changes in the kidney of diabetic rats. Treatment with *F. deltoidea* ameliorates kidney injury in STZ-treated rats, at least in part, by ameliorating hyperglycemia-mediated oxidative stress and apoptosis. We also provide the first evidence that there is a good agreement between the intensity of IR peaks at 1545 cm^−1^, 1511 cm^−1^ and 1246 cm^−1^ and kidney function, supporting the use of FTIR in accessing the nephroprotective effect of *F. deltoidea*.

## Acknowledgments

This research was supported by grants from the Ministry of Science, Technology and Innovation (SF: 100-RMI/SF 16/6/2 (7/2015), Ministry of Higher Education (MOE FRGS: 600-RMI/FRGS 5/3 (5/2014), Faculty of Applied Sciences, Universiti Teknologi MARA (UiTM). We thank Laboratory Animal Facility and Management (LAFAM), UiTM for the postoperative care of the animals.

## Author contributions statement

Nurdiana Samsulrizal, designed and carried out the studies, performed the statistical analysis, interpreted the results, and drafted the manuscript. Goh Yong-Meng. supervised, designed the experiments, and drafted the manuscript. Hafandi Ahmad participated in biochemical analysis. Nur Syimal’ain Azmi and Noor Syaffinaz Noor Mohamad Zin carried out the animal handling. Ebrahimi Mahdi performed the biochemical and statistical analyses. All authors reviewed the manuscript.

